# A new sequencing-based women’s health assay combining self-sampling, HPV detection and genotyping, STI detection, and vaginal microbiome analysis

**DOI:** 10.1101/217216

**Authors:** Elisabeth M. Bik, Sara W. Bird, Juan P. Bustamante, Luis E. Leon, Pamela A. Nieto, Kwasi Addae, Víctor Alegría-Mera, Cristian Bravo, Denisse Bravo, Juan P. Cardenas, Adam Caughey, Paulo C. Covarrubias, José Pérez-Donoso, Graham Gass, Sarah L. Gupta, Kira Harman, Donna Marie B. Hongo, Juan C. Jiménez, Laurens Kraal, Felipe Melis-Arcos, Eduardo H. Morales, Amanda Morton, Camila F. Navas, Harold Nuñez, Eduardo Olivares, Nicolás Órdenes-Aenishanslins, Francisco J. Ossandon, Richard Phan, Raul Pino, Katia Soto-Liebe, Ignacio Varas, Patricia Vera-Wolf, Nathaniel A. Walton, Daniel E. Almonacid, Audrey D. Goddard, Juan A. Ugalde, Jessica Richman, Zachary S. Apte

## Abstract

The composition of the vaginal microbiome, including both the presence of pathogens involved in sexually transmitted infections (STI) as well as commensal microbiota, has been shown to have important associations for a woman’s reproductive and general health. Currently, healthcare providers cannot offer comprehensive vaginal microbiome screening, but are limited to the detection of individual pathogens, such as high-risk human papillomavirus (hrHPV), the predominant cause of cervical cancer. There is no single test on the market that combines HPV, STI, and microbiome screening. Here, we describe a novel inclusive women’s health assay that combines self-sampling with sequencing-based HPV detection and genotyping, vaginal microbiome analysis, and STI-associated pathogen detection. The assay includes genotyping and detection of 14 hrHPV types, 5 low-risk HPV types (lrHPV), as well as the relative abundance of 32 bacterial taxa of clinical importance, including *Lactobacillus*, *Sneathia*, *Gardnerella*, and 4 pathogens involved in STI, with high sensitivity, specificity, and reproducibility. For each of these taxa, healthy ranges were determined in a group of 50 self-reported healthy women. The hrHPV portion of the test was evaluated against the Digene High-Risk HPV HC2 DNA test with vaginal samples obtained from 185 women. Results were concordant for 181/185 of the samples (overall agreement of 97.83%, Cohen’s kappa = 0.93), with sensitivity and specificity values of 94.74% and 98.64%, respectively. Two discrepancies were caused by the Digene assay’s known cross-reactivity with low-risk HPV types, while two additional samples were found to contain hrHPV not detected by Digene. This novel assay could be used to complement conventional cervical cancer screening, because its self-sampling format can expand access among women who would otherwise not participate, and because of its additional information about the composition of the vaginal microbiome and the presence of pathogens.

## Introduction

A woman’s vaginal health is critical for her general well-being and reproductive success, and is determined by the vaginal microbiome composition, the presence of pathogens associated with sexually transmitted infections (STI), and the presence of human papillomavirus (HPV) types that can cause genital warts or cervical cancer. Current clinical women’s health assays focus on the detection of STI or that of HPV, but there is no single test that combines these targets with vaginal microbiome analysis or with self-sampling.

The human vaginal microbiome has a unique composition compared to other microbial communities in the human body. In most healthy women, the vaginal microbiome is characterized by a low bacterial diversity and a dominance of lactobacilli, with low abundance of other bacterial genera (Ravel *et al.* 2011; Younes et al. 2017). Several vaginal microbial community types have been described, most of which are dominated by a single *Lactobacillus* species. Lactic acid produced by lactobacilli lowers the vaginal pH, which is believed to create an environment unfavorable for the growth of pathogenic bacteria (O’Hanlon et al. 2013). Low numbers of vaginal lactobacilli have been associated with many health conditions, such as bacterial vaginosis (Ling et al. 2010; Ravel et al. 2011; Ravel et al. 2013; Srinivasan et al. 2012), aerobic vaginitis (Donders et al. 2017), cervicitis (Gorgos et al. 2015), and STI (Petrova et al. 2015; Hill et al. 2016; Jensen 2017; Ziklo et al. 2016). The composition of a woman’s vaginal microbiome thus plays an important role in women’s health and reproductive success. Yet, the analysis of this microbial community is not part of regular health care for women. In the US, as in many other countries, most healthcare providers instead focus on the detection of high-risk human papillomavirus (hrHPV), the predominant cause for cervical cancer.

Cervical cancer is one of the major causes of cancer-related deaths in women, with an annual worldwide mortality of 250,000 (Bray et al. 2013; Jemal et al. 2011). hrHPV DNA can be detected in almost all (>99%) cervical cancer specimens, and HPV is therefore considered the predominant causative agent for cervical cancer (Bosch and Muñoz 2002; Walboomers et al. 1999). Although HPV infection is the most common STI worldwide, not all HPV infections will lead to cancer. Firstly, certain HPV types have higher oncogenic risks than others. Of the over 170 different HPV genotypes known to date, twelve types have been classified as Group 1 human carcinogens; these include types 16, 18, 31, 33, 35, 39, 45, 51, 52, 56, 58, and 59 (Bouvard et al. 2009; IARC Working Group on the Evaluation of Carcinogenic Risks to Humans 2012). Together with other, closely related HPV types such as 66 and 68, which have been listed as probably or possibly carcinogenic, these are collectively called high risk HPV (hrHPV) types. hrHPV types 16 and 18 can be found in over 70% of cervical cancers (de Sanjose et al. 2010) and the presence of these types is associated with the highest chance of developing cancer within 10 years (Khan et al. 2005). However, other hrHPV genotypes have also been shown to cause cervical cancer, and especially among women of non-European descent (Vidal et al. 2014). In addition, many hrHPV infections are temporary and will be cleared within months of acquisition, without proceeding to pre-cancerous lesions (Rosa et al. 2008). Other HPV types, collectively called low-risk HPV (lrHPV), are not implicated in cervical cancer, but instead cause genital warts (Egawa and Doorbar 2017).

National cervical cancer screening programs are offered to women over 21 years old worldwide (Mendes et al. 2015). Most of these programs involve an invitation for a Pap smear, in which a woman’s cervical cells are obtained by a physician for cytology (Tambouret 2013), but additional molecular HPV testing is increasingly offered by health care providers as well (Tota et al. 2017). Studies suggested that clinical tests based on detection of HPV DNA exhibit higher sensitivity for detection of cervical intraepithelial neoplasia, in comparison with Pap testing (e.g. Mayrand et al. 2007). In the United States, most healthcare providers follow the American College of Obstetricians and Gynecologists (ACOG) guidelines (Committee on Practice Bulletins—Gynecology 2016) or the U.S. Preventive Services Task Force (USPSTF) guidelines for women (US Preventive Services Task Force 2016) to come in for a Pap smear, often with HPV testing, every three to five years, depending on age and risk factors.

Several commercial kits have FDA pre-market approval for the molecular detection of HPV (Gradíssimo and Burk 2017). In a 2014 meta-analysis of 36 studies, the Qiagen Digene^®^ Hybrid Capture^®^ 2 (HC2) assay was the most widely used (Arbyn et al. 2014). In the HC2 assay (de Roda Husman et al. 1995), vaginal specimens are denatured with sodium hydroxide, denatured viral DNA is hybridized with specific RNA probes, and RNA:DNA hybrids are subsequently detected with antibodies (Lörincz 1996). The HC2 test detects 13 hrHPV types, but does not report which specific type is present. Other HPV detection assays involve the amplification of viral DNA by Polymerase Chain Reaction (PCR). The most widely used primer pairs for HPV PCR detection are the GP5+/6+ primers (de Roda Husman et al. 1995) and the degenerate MY09/11 primers (Manos et al. 1989) or the PGMY09/11 primer pool (Gravitt et al. 2000), which are all based on conserved regions in the viral L1 open reading frame. The COBAS^®^ 4800 assay detects 14 hrHPV types using multiplex real-time PCR with specific probes; it reports the presence of HPV16, HPV18, or one of 12 remaining hrHPV types (Heideman et al. 2011).

Despite the offering of cervical cancer screening services, only 81% of women in the US participate in cervical cancer screening programs (National Center for Health Statistics. 2016; Watson et al. 2017). About one in five US women aged 21–65, a group of 14 million women, have not been screened in the past 3 years (Watson et al. 2017). Screening participation is particularly low among certain populations such as American Indians, Asians, Native Hawaiians, and recent immigrants, as well as women who live below poverty level or who have experienced partner violence (Musselwhite et al. 2016; National Center for Health Statistics. 2016; Levinson et al. 2016; Watson et al. 2017). Failure to respond to cervical cancer screening invitations and reminders can be caused by a combination of different factors. The most important reasons for attendance failure are lack of time for a visit to a clinic, embarrassment to undergo a pelvic exam, and memories of discomfort or pain at previous clinical visits (Dzuba et al. 2002; Sultana et al. 2015).

In the US, self-screening for HPV testing is not yet recommended as part of the standard of care. However, a number of countries have already switched to or are considering to offer self-sampling and HPV testing as a way to increase attendance for cervical cancer screening. In 2016, the Netherlands was the first country to start a new screening program that allows women to self-collect samples for HPV testing (Rozemeijer et al. 2015). Clinical trials in many other countries are ongoing, including Australia (Sultana et al. 2016), Denmark (Tranberg et al. 2016), Finland (Virtanen et al. 2015), Italy (Giorgi Rossi et al. 2015), Norway (Enerly et al. 2016), Switzerland (Viviano et al. 2017), and the UK (Lim et al. 2017). Offering women the opportunity to self-collect vaginal specimens poses fewer barriers for women to be screened, leading to increased participation rates (Verdoodt et al. 2015). Thus, encouraging women to self-collect vaginal samples for hrHPV screening may have an impact on rates of detection of cervical cancer (Wong et al. 2016).

The vaginal microbiome is an emerging area of research in understanding the role of HPV infections and reducing the risk of cervical cancer (Mitra et al. 2016). Several studies suggest a relationship between the composition of the vaginal microbiota and the acquisition and persistence of HPV infection. For example, vaginal microbial diversity is increased during an HPV infection, with decreased levels of *Lactobacillus* species and an increased presence of other microbial members such as *Sneathia* species or *Gardnerella vaginalis* (Gao et al. 2013; Lee et al. 2013; Brotman et al. 2014; Reimers et al. 2016; Shannon et al. 2017). In addition, certain microbiota compositions are associated with increased clearance of detectable HPV (Brotman et al. 2014).

In this study, we tested the feasibility of a novel assay, that combines the detection and identification of HPV DNA, STI-associated pathogens, and microbiome analysis on samples obtained through self-sampling. We validated the performance of marker gene amplification and sequencing to detect the presence and relative abundance of 32 clinically important bacterial targets with high precision and accuracy. In addition to detecting *Lactobacillus*, *Sneathia*, and *Gardnerella* spp., this test detects STI-associated pathogens including *Chlamydia trachomatis, Mycoplasma genitalium, Neisseria gonorrhoeae,* and *Treponema pallidum,* which cause chlamydia, genital tract infections, gonorrhea, and syphilis, respectively. The performance of a novel amplification and sequencing-based strategy for HPV detection and type-specific identification was compared to that of the most widely used test for HPV detection in cervicovaginal specimens, the Digene HC2 test. This assay is intended to complement, rather than replace, current healthcare guidelines for in-clinic cervical cancer screening.

## Materials and methods

### Study participants and sample collection

The specimens used in this study consisted of vaginal samples from women who had signed an informed consent to have their samples used for research. This study was approved under a Human Subjects Protocol provided by an IRB (E&I Review Services, IRB Study #13044, 05/10/2013). All participants were 18 years or older. A vaginal self-collection kit was sent to each participant’s home address, consisting of a sterile swab, a tube with sterile water, a tube with zirconia beads in a proprietary lysis and stabilization buffer that preserves the DNA for transport at ambient temperatures, and sampling instructions (Supplementary Figure 1, included in the Supplementary Materials). Participants were instructed to wet the swab with the sterile water, insert the swab into the vagina as far as is comfortable, make circular movements around the swab’s axis for 1 minute (min), and then stir the swab for 1 min into the tube with lysis buffer and beads. After shaking the tube for 1 min to homogenize, the tube was then shipped by the participants to the laboratory by regular mail.

For the determination of the healthy ranges of the 32 bacterial targets, a set of 50 vaginal specimens, each from a different woman (average age 48.4 ± 15.6 years), was selected. Inclusion criteria were the following: completion of the voluntary health survey that every woman was invited to participate in, and no report of the following conditions: bacterial vaginosis, cervical cancer, genital herpes or warts, urinary tract infection, or infection with HPV, *C. trachomatis, T. pallidum*, or yeast. In addition, all of these women reported no antibiotic usage in the six months before sampling.

A different set of specimens from 88 women was used to compare the performance of sampling with the Digene collection device (Qiagen, Gaithersburg, MD, USA) and DNA extracted from samples collected with swabs. For this subset, women were asked to self-sample 2 vaginal specimens within 15 minutes. The first specimen was collected by using the Digene collection device, which consists of a cervical brush and a Digene transport tube with Specimen Transport Medium (STM). The second specimen was collected using a pre-wetted swab and resuspended in a collection tube with lysis buffer and beads, as described above, and used for DNA extraction.

A third set of vaginal specimens from 185 women was selected to compare the performance of the Digene HC2 HPV assay versus that of the amplification and sequence-based HPV type identification described in this study.

For use in some experiments described below, homogenized “vaginal pools” were created by combining 96 vaginal samples derived from 11 or 16 individuals who sampled themselves multiple times.

### Positive STI control samples

Ten de-identified cervicovaginal swab specimens of known STI pathogen status were obtained through a commercial source (iSpecimen, Lexington, MA). Five of these samples were reported to be positive for *C. trachomatis* and negative for *N. gonorrhoeae*, while a second set of five samples were negative for *C. trachomatis* and positive for *N. gonorrhoeae.* Each sample was tested in five replicates for DNA extraction, 16S rRNA gene amplification, and target identification as described below.

### *In silico* 16S rRNA gene target performance metrics

The assay includes 32 bacterial targets with clinical relevance for women’s reproductive tract health, which were identified through an exhaustive literature search (Figure 1). These targets were chosen based on a review of the clinical and scientific literature related to vaginal health. The most relevant associations between health conditions and the vaginal microbiota were narrowed down by choosing associations with high statistical significance that were found in humans subjects, not in laboratory animals or bioreactors, but performed on case/control, cohorts or randomized studied population. These include bacterial vaginosis (Ling 2010, Ravel 2011, Ravel 2014, Srinivasan 2012), aerobic vaginitis (Donders 2017), pelvic inflammatory disease (Brunham 2015), and sexually transmitted infections (Hill 2016; Jensen 2017; Ziklo 2016). A complete list of these associations and references are provided in Supplementary Table 1. For each bacterial taxon intended to be included in this assay, using a process similar to that described in Almonacid et al. (2017), we determined *in silico* performance metrics for identification of each taxa (sensitivity, specificity, positive and negative predictive value). Briefly, sequences assigned to each taxa in the Silva database (Version 123) (Quast, 2013) were considered to be real positives for that taxa.

**Figure 1.**
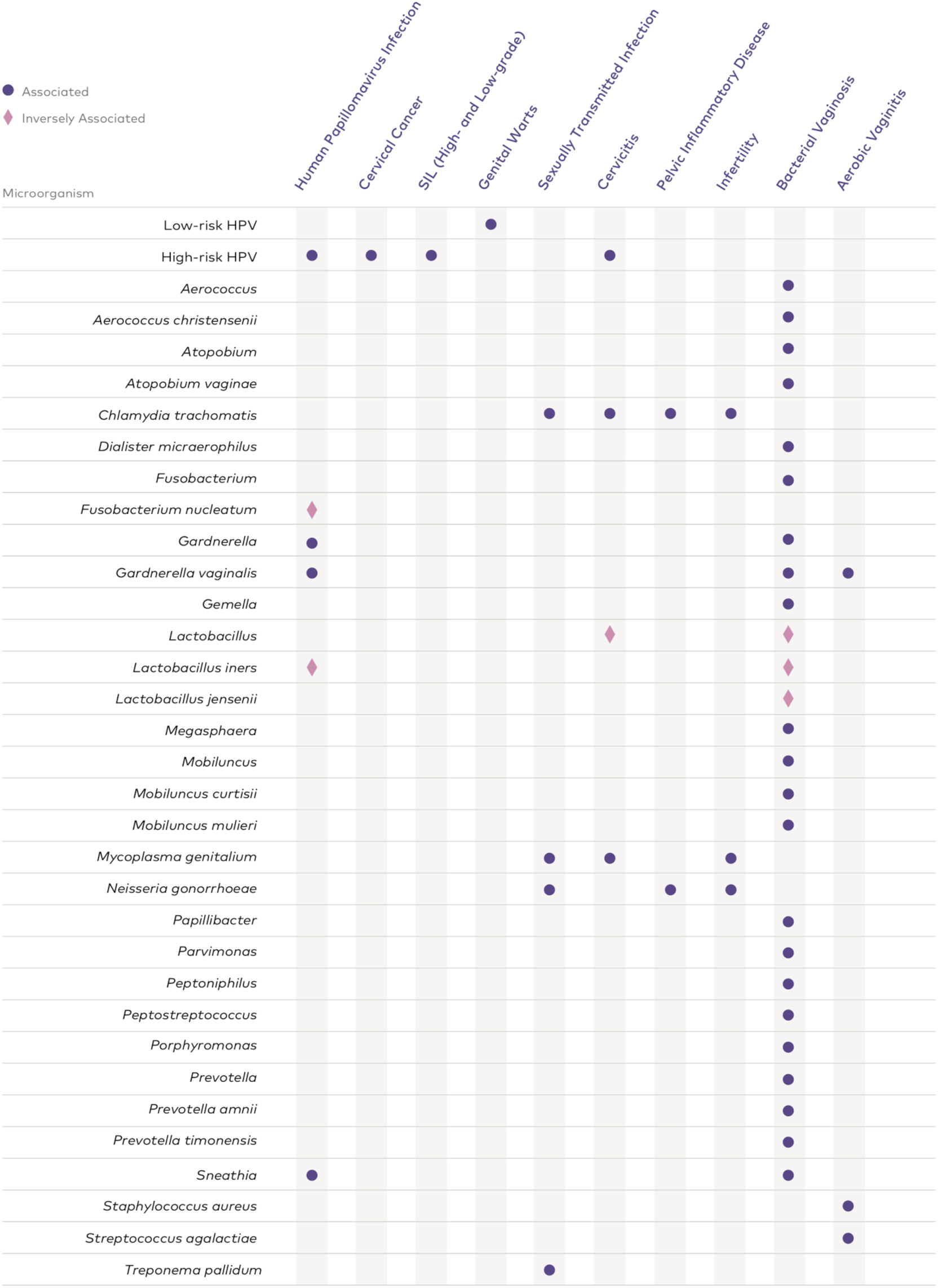
**The 32 bacterial targets and HPV targets covered by the assay and their associated health conditions.** See Supplementary Table 1 in the Supplementary Materials for more detailed information about e.g. the HPV genotypes included and a list of references.

Then, assuming amplification with up to two mismatches with the primers used, we identified for each taxa the sequences that would produce an amplicon, and evaluated whether that amplicon is unique to the taxon of interest (ti) or also shared by sequences from different taxa (dt). The number of true positives (TP), true negatives (TN), false positives (FP) and false negatives (FN) was computed for different tolerance ratios for the quotient dt/ti, and subsequently *in silico* performance metrics were assessed. Of the 73 bacterial targets initially selected, the 32 targets selected for the assay had all four *in silico* performance metrics above 90% (Supplementary Figure 2).

### *In silico* HPV target performance metrics

In addition to the 32 bacterial targets, hrHPV and lrHPV targets were selected for inclusion in the assay, based on their association with cervical cancer lesions or genital warts (Figure 1, Supplementary Table 1). HPV reference genomes were downloaded in August 2017 from the PaVE database, which is a repository of curated and annotated HPV genomes (Van Doorslaer et al. 2013, Van Doorslaer et al. 2017). Only revised and recognized sequences (180 HPV genomes) were used for an *in silico* PCR amplification using a set of 15 forward and 6 reverse primers (described below) targeting the L1 gene and allowing up to 4 mismatches between primers and target sequences. Under these conditions, the L1 genes from 118 HPV genomes could be amplified *in silico*. Of these, 19 HPV genomes, including 14 hrHPV types (i.e., types 16, 18, 31, 33, 35, 39, 45, 51, 52, 53, 56, 58, 59, 66, 68) and 5 lrHPV types (6, 11, 42, 43, 44) were selected based on their association with health conditions according to literature (Supplementary Table 1). In order to evaluate the performance metrics for identification of the HPV targets, sequences of the L1 segment of HPV genomes from the NCBI database were used. The search was filtered to sequences with length in the range 1,500-10,000 bp and with correct assignment of the type of the HPV (4177 sequences). These sequences were amplified *in silico* using the primers described below. Following these steps, we generated 161,398 amplicons. These sequences were mapped using Vsearch (Rognes 2016) at 95% of identity against an HPV amplicon reference database consisting of the amplicons produced by the reference genomes in PaVe for the 19 HPV types included in our assay. The performance metrics were calculated in a manner analogous to how they were calculated for the 16S rRNA gene targets. Briefly, the correct assignment of an NCBI amplicon against the reference was counted as a true positive and an incorrect assignment was considered as a false negative. Also, we considered as a false negatives the genomes from NCBI that our primers could not amplify. According to this, the 19 HPV types obtained values for sensitivity, specificity, positive predictive value (PPV) and negative predictive values (NPV) above 90% (Supplementary Table 2).

### *In vitro* validation of bacterial targets

To test our ability to identify members of each of the 32 bacterial targets, synthetic DNAs (sDNAs or gBlocks, Integrated DNA Technologies, Inc., Coralville, IA), were designed encompassing the V4 region of the 16S rRNA gene including primer regions, based on a Silva representative sequence, plus 75 additional bases to both the 5’ and 3’ side, with one sDNA per target (Supplementary Table 3). The Silva representative sequence per taxa was chosen by performing an all-against-all sequence comparison of all sequences in a taxa, and identifying as representative the sequence that shared the highest similarity with the largest number of sequences in the set.

To validate that each target could be detected in a vaginal swab specimen, 3 ng of each sDNA was spiked into 500 µl aliquots of a vaginal pool, created by combining 96 vaginal specimens of women included in this study, and DNA was extracted from each spiked vaginal pool (see below). Each spike-in experiment was performed in triplicate. Subsequently, bacterial targets were detected by amplification using PCR targeting the 16S rRNA gene, sequencing, and a bioinformatics pipeline described below. Each target included in the final panel was detected above limit of detection (LOD) (see below) in each of the triplicate spiked-in amplification reactions performed on the extracted DNA from the vaginal pool (not shown).

In addition, the LOD of each target in a complex background of other targets was determined according to published guidelines (Clinical and Laboratory Standards Institute 2004). First, we calculated a limit of blank (LOB), which was calculated using a set of 77 blank wells of a 96-well PCR plate where wells of the first row and first column of the plate each contained 200 pg/µl of synthetic 16S rRNA gene DNA from different targets. The LOB was set as the average number of reads in these blank wells (18.57 reads) plus 1.65 standard deviations (29.70 reads), thus at 48.27 reads. To calculate the LOD of the bacterial target, pools of bacterial sDNAs were mixed in different ratios. To create these mixes, each bacterial sDNA was randomly assigned to one of two pools, A and B, that each contained sDNAs in equimolar amount. Each pool was serially diluted in PCR grade water. Pool A dilutions were mixed 1:1 with undiluted Pool B and vice versa. All pool A/B combinations were used in triplicate for DNA extraction, amplification, and sequencing as described below. For each target, the LOD was defined as the lowest concentration of sDNA where at least two of the three replicates contained at least 2 reads for that target in a sample with 10,000 reads or more. Using this LOD, we calculated a lower threshold for detection for each taxa at its LOD as the LOB (48.27) plus the standard deviation of the taxa at LOD * 1.65. This threshold is used to correctly assign a taxa as identified in a sample at or above its LOD.

For targets that had both a species and a genus level sDNA present in the mixed pools A and B, a bioinformatic correction was applied. The total reads for a genus-level target for which a species within that genus was also present in the mixed pools, was defined as the total measured reads for the genus and subtracting all those reads corresponding to species-level targets belonging to that genus in the same pool mix, i.e., only reads that match to a genus and not to a species level were finally assigned to the genus.

### *In vitro* validation of HPV targets

To test the ability of our assay to detect and genotype HPV targets, fragments of the L1 gene of approximately 600bp long were ordered for each of the representative sequences of 19 HPV types in the PaVE database as sDNAs (gBlocks, Integrated DNA Technologies, Inc.). To represent hrHPV type 68, two sDNAs were ordered, 68a and 68b. The sequences of the 20 gBlocks representing 19 HPV types (14 hrHPV and 5 lrHPV) are listed in Supplementary Table 4. To validate that each target could be detected in a vaginal swab specimen, 3 ng of each HPV sDNA was spiked into 500 µl aliquots of a vaginal pool created by combining 96 vaginal specimens of women included in this study, and DNA was extracted from each spiked vaginal pool (see below). Subsequently, the spiked HPV targets were detected by amplification using the PCR targeting the L1 gene and bioinformatics pipeline described below. Each spike-in experiment was performed in triplicate. Each HPV target was detected above the LOD (see below) in each of the triplicate spiked-in amplification reactions performed on the extracted DNA from the vaginal pool (not shown). Each target had a ratio > 0.1 for the number of HPV-assigned reads divided by the total number of normalized reads assigned to an internal spike-in control (see below).

To determine the LOD of HPV targets, 10-fold serial dilutions of the sDNAs representing HPV targets were made in nuclease-free water, ranging from 10^5^ to 10^2^ molecules per µl. Dilutions of one target were inversely combined with dilutions of another target, forming different pairs of HPV sDNAs. Each dilution pair was used directly as template for PCR in triplicate as described below.

### DNA extraction and amplification targeting 16S rRNA and HPV L1 genes

DNA was extracted from vaginal specimens, pools thereof, or sDNA dilutions in tubes containing lysis/stabilization buffer as described previously (Almonacid et al. 2017). For 16S rRNA gene amplification, extracted DNA was used as the input of a one-step PCR protocol to amplify the V4 variable region of the 16S rRNA gene. This PCR contained universal primers 515F and 806R (Almonacid et al. 2017; Caporaso et al. 2011), both with sample-specific indices and Illumina tags. PCR was performed as described before (Almonacid et al 2017). Following amplification, DNA was pooled by taking the same volume from each reaction.

For HPV amplification, extracted DNA was used as the input of a PCR protocol to amplify the HPV L1 gene. To each sample, sDNA with a randomized HPV type 16 sequence was added as an internal positive control. The first PCR mix contained a pool of previously described HPV specific primers (Gravitt et al. 2000; Estrade and Sahli 2014), and two new primers, HPV_RSMY09-LvJJ_Forward: 5’ CGTCCTAAAGGGAATTGATC, and HPV_PGMY11-CvJJ_Reverse: 5’ CACAAGGCCATAATAATGG. All these primers contained sequencing adaptor regions. The PCR products from the first amplification round were used as input for a second PCR containing sample-specific forward and reverse indices and Illumina tags. PCR products from this second step were pooled for sequencing.

The 16S rRNA gene and HPV PCR consolidated library pools were separately quantified by qPCR using the KAPA Library Quant Kit (Bio-Rad iCycler qPCR Mix) following the manufacturer’s instructions using a BioRad MyiQ iCycler. Sequencing was performed in a paired-end modality on the Illumina NextSeq 500 platform rendering 2 x 150 bp pair-end sequences.

### Sequence analysis and taxonomic annotation for bacterial targets

After sequencing, demultiplexing of reads according to sample-specific barcodes was performed using Illumina’s BCL2FASTQ algorithm. Reads were filtered using an average Q-score > 30. Forward and reverse 16S rRNA gene reads were appended together after removal of primers and any leading bases, and clustered using version 2.1.5 of the Swarm algorithm (Mahe 2014) using a distance of one nucleotide and the “fastidious” and “usearch-abundance” flags. The most abundant sequence per cluster was considered the real biological sequence and was assigned the count of all reads in the cluster. The representative reads from all clusters were subjected to chimera removal using the VSEARCH algorithm (Rognes 2016). Reads passing all above filters (filtered reads) were aligned using 100% identity over 100% of the length against the 32 target 16S rRNA gene sequences described above (Supplementary Table 4). The relative abundance of each taxon was determined by dividing the count linked to that taxa by the total number of filtered reads.

### Sequence analysis and taxonomic annotation for HPV targets

Raw sequencing reads were demultiplexed using BCL2FASTQ. Primers were removed using cutadapt (Martin 2011). Trimmomatic (Bolger 2014) was used to remove reads with a length less than 125 bp, and a mean quality score below 30. After that, forward and reverse paired reads were joined using custom in-house scripts and converted to a fasta file. Identical sequences were merged and written to a file in fasta format and sorted by decreasing abundance using --derep_fulllength option in VSEARCH (Rognes 2016). Target sequences in the fasta files were compared to the fasta-formatted query database sequences (19 HPV target sequences) using the global pairwise alignment option with VSEARCH, using 95 percent sequence identity, to obtain the counts for each HPV type within a different sample.

The HPV portion of the assay was considered positive if the number of sequence reads assigned to the specific HPV types was above the threshold at the limit of detection, and greater than a previously defined cutoff. To set this cutoff, two normalization steps were employed. First, according to *in silico* PCR amplification, a different number of combinations of primers amplify different HPV targets (e.g. HPV16 is amplified using 66 different combinations, while HPV43 is amplified with just 10 combinations), reflecting the sequence variability within the primer binding site among HPVs. This also means that the spiked-in internal control and the target HPV have different amplification efficiencies. To avoid this bias, the internal control (which has the primer sites for HPV16) is normalized for the amplification factor (number of primer combinations that generate an amplicon) of each HPV type. The number of HPV-assigned reads was divided by the total number of normalized reads assigned to the spike, and a sample was considered HPV-positive if that ratio was above 0.1, which was obtained from the concordance study, and corresponds to approximately 500 target molecules.

### Intra- and inter-run precision

Intra-run technical repeatability was assessed by including nine replicates of the same vaginal pool (consisting of 96 vaginal samples derived from 11 individuals) into the same DNA extraction, 16S rRNA gene amplification, and sequencing run. This experiment was then repeated in a second sequencing run to yield another set of nine replicate samples analyzed within the same run. In addition, inter-run technical reproducibility was performed by processing three replicates of a set of 18 vaginal samples on three different days by three different operators. Samples included in the analysis were those that had at least 10,000 reads and where at least two of the three replicates were present (11 sets).

Comparison of the results, both intra- and inter-run, were done using the raw counts of the 32 bacterial species- and genus-level targets. Data was processed using the R-package Phyloseq (McMurdie and Holmes 2013), visualized using Principal Coordinates Analysis (PCoA), based on a distance matrix calculated using the Bray-Curtis method.

### Digene HC2 hrHPV test on Digene tubes or on extracted DNA

The Digene HC2 High-Risk HPV detection assay (Qiagen) was used as a reference to validate the hrHPV portion of the assay. The High-Risk HPV Probe in the Digene HC2 HPV test detects hrHPV types 16, 18, 31, 33, 35, 39, 45, 51, 52, 56, 58, 59, and 68, while the women’s health test described here detects all these types plus HPV66. The Digene HC2 assay is intended to be used directly on vaginal samples collected in the Digene STM transport tube. In order to validate the use of the Digene kit on extracted vaginal DNA, we compared the performance of the Digene HC2 assay on a set of 88 self-obtained, paired vaginal samples, i.e. a Digene brush resuspended in STM, as well as DNA extracted from a vaginal swab resuspended in lysis and stabilization buffer. For the specimens collected in STM, 500 µl of specimen sample was mixed with 250 µl of Denaturing Reagent (provided in the Digene HC2 kit), shaken on a Hybrid Capture System Multi-Specimen Tube Vortexer (Digene) for 30 seconds, and incubated in a waterbath at 65°C for 45 min, as per the manufacturer’s instructions. Alternatively, for specimens collected with the women’s health assay kit, 50 µl of extracted DNA was mixed with 25 µl of denaturing reagent and incubated at 65°C for 45 min in the Microplate Heater I (Digene). For both sample types, denatured samples were then hybridized to the High-Risk HPV RNA probe set in the Microplate Heater I at 65°C for 60 min, and DNA:RNA hybrid molecules were detected using monoclonal antibodies and chemiluminescence measurement on a DML3000 machine, according to the manufacturer’s instructions (Digene^®^ HC2 HPV DNA Test Instructions For Use, Qiagen). Negative and positive controls and calibrators included in the kit were processed within each 96-well assay, and used for assay validation and cutoff, as per instructions. A specimen was considered positive if its chemiluminescence measurement (Relative Light Units, RLU) was higher than or equal to that of the assay’s Positive Calibrator cutoff (RLU ratio of 1 or more), as specified in the Digene HC2 assay instructions.

Sensitivity, specificity, and accuracy of the hrHPV portion of the women’s health assay were evaluated using the Digene HC2 hrHPV assay as the gold standard and extracted DNA from 185 vaginal swabs as the input for both tests. The assay was considered to be positive for hrHPV if the number of reads assigned to hrHPV types divided by the normalized number of reads assigned to a spiked-in control (see above) was greater than 0.1. For this comparison, sequences assigned to hrHPV type 66 were not considered, because this type is not detected in the Digene assay. Agreement between the two methods was evaluated using Cohen’s kappa (Cohen 1960), where the level of agreement is defined by the range: 0-0.2, poor; 0.21-0.40, fair; 0.41-0.6, moderate; 0.61-0.8, good; 0.81-1.00, very good.

## Results

### Limit of detection of bacterial and HPV targets

The test is based on a list of 32 bacterial 16S rRNA gene targets and 19 HPV types that were identified through an exhaustive literature search to play important roles in health and disease of women’s reproductive tracts (Figure 1, Supplementary Table 1). For each bacterial target, the LOD was determined by combining different dilutions of pools of sDNAs, followed by DNA extraction, amplification of the V4 region of the 16S rRNA gene using broad range primers, and sequencing (Figure 2). The LOB was set as the average number of reads in 77 blank wells (18.57 reads) plus 1.65 standard deviations (29.70 reads). Using this value, we calculated the threshold of identification for each taxon as the LOB + 1.65 standard deviations (48.27) plus the standard deviation of the taxon at LOD * 1.65 (Supplementary Table 5). For example, the LOD for *Atopobium vaginae* was identified at the 1:100 dilution, and its corresponding threshold was set at 49 reads. For the other 31 taxa targeted by the assay, the threshold related to LODs was in the range 49.0 to 65.2 reads (Supplementary Table 5).

**Figure 2.**
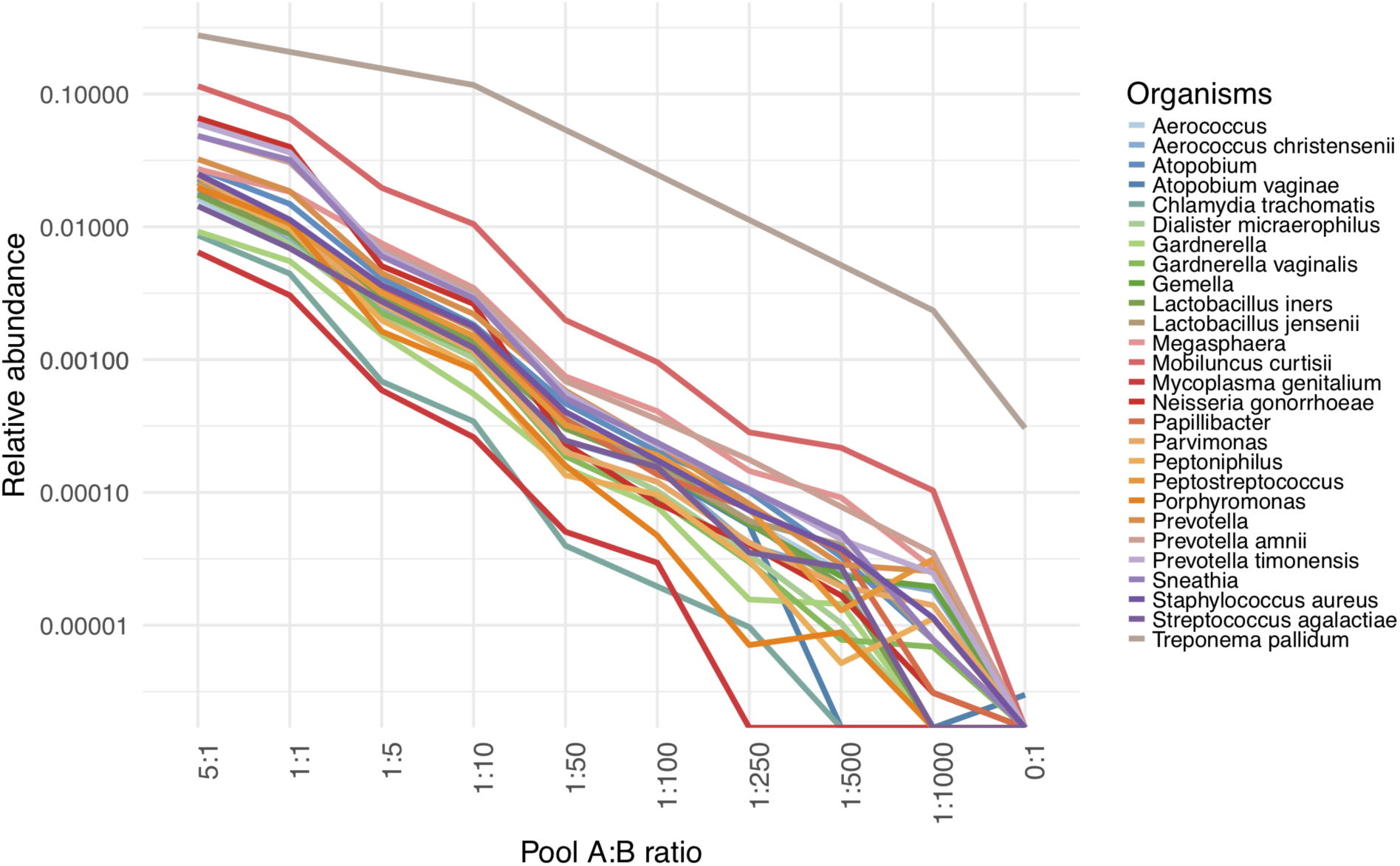
**Limit of detection of sDNAs representing bacterial 16S rRNA gene sequences in a complex background of other sDNAs.** Dilutions of two pools of sDNAs were mixed in different amounts, and bacterial targets were amplified and sequenced. For each dilution and target, the relative abundance in samples with 10,000 reads or more are shown. Unlike all other targets, which were mixed in pools consisting of 30 targets, the *T. pallidum* sDNA was tested in pools containing only 2 targets; hence its relative abundance in the mixed pools was much higher than that of the other targets, explaining why the *T. pallidum* curve is shifted to the right. LOD read thresholds are provided in Supplementary Table 5.

To determine the LOD for the HPV targets, different dilutions of pools of sDNAs were mixed as done for the bacterial targets. The molecules were then amplified, sequenced, and analyzed by the HPV bioinformatics pipeline. For all HPV targets analyzed, the threshold related to LODs was in the range 40.8 to 224.8 reads (Figure 3, Supplementary Table 6).

**Figure 3.**
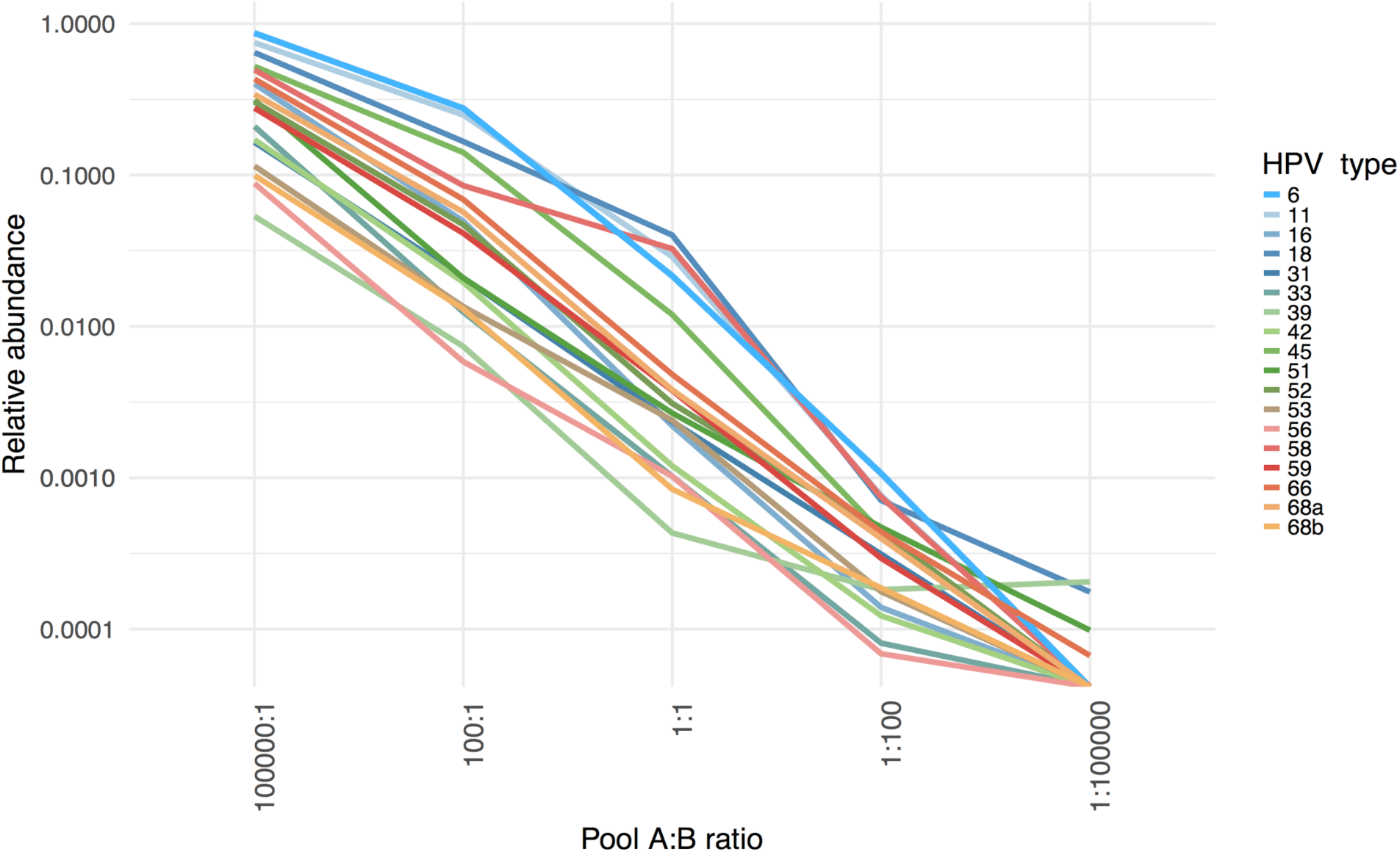
**Limit of detection of HPV types**. Dilutions of two pools of sDNAs representing HPV types were mixed in different amounts, and HPV types were amplified and sequenced. Two different sDNAs were used to represent hrHPV type 68. For each dilution and HPV type, the relative abundance in samples with 10,000 reads or more are shown. The LOD read thresholds for each HPV target are provided in Supplementary Table S7.

### Intra- and inter-run variability

Intra-run technical variability was evaluated in a combined set of 18 replicates of the same vaginal pool, each of which yielded 10,000 reads or more. Ordination plots of both genus and species level bacterial communities (Figure 4) showed a tight clustering of intra-run technical replicates, indicating that within a single sequencing run, results generated by the laboratory process and the bioinformatics analysis were consistent.

**Figure 4.**
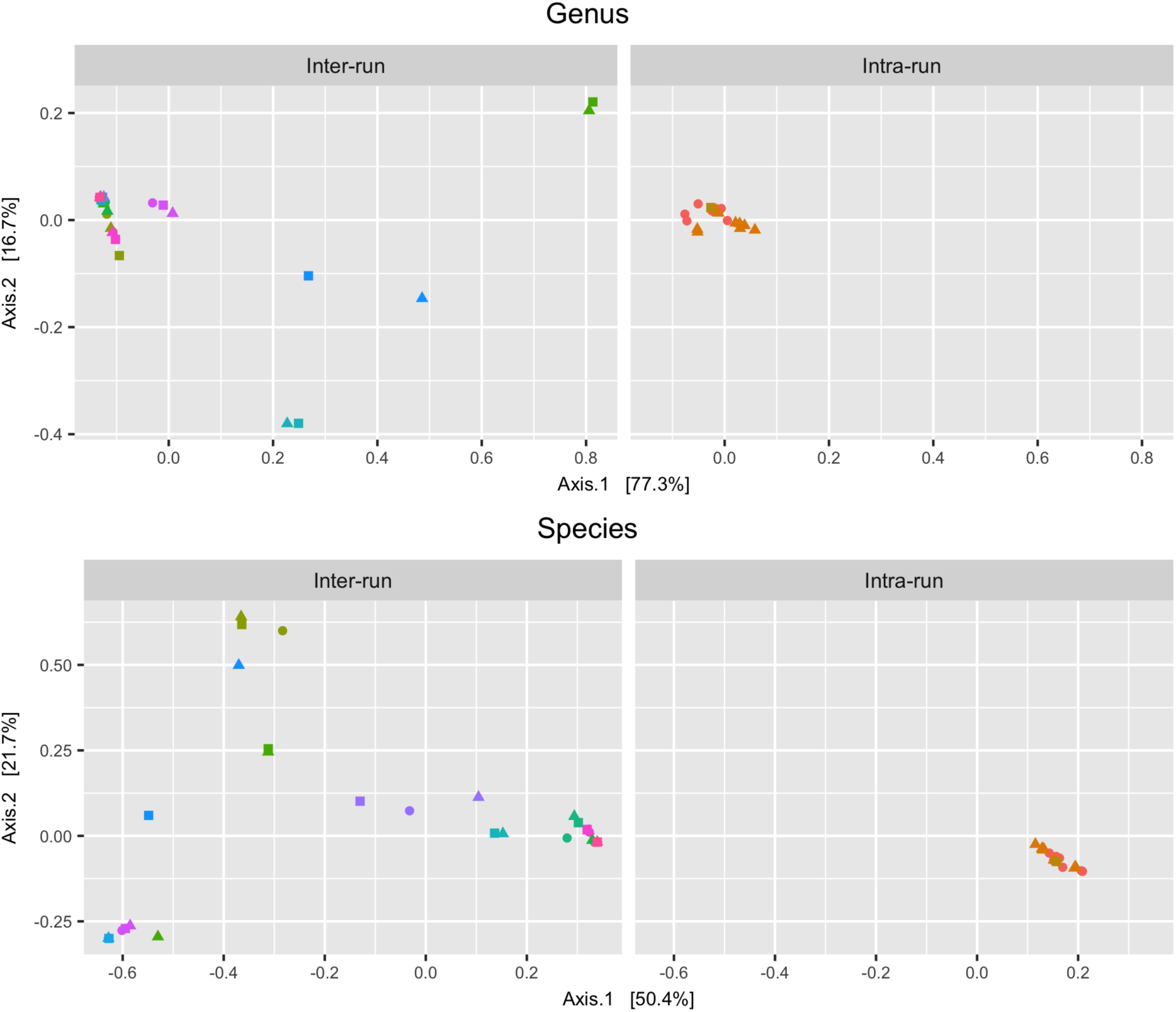
**Inter- and intra-run reproducibility.** PCoA ordination showing clustering of inter-run (11 vaginal samples analyzed in triplicate on three independent sequencing runs) and intra-run (18 aliquots of the same vaginal pool) data, at the genus and species taxonomic level. Shapes indicate the sequencing run, while colors indicate sample replicates.

For the inter-run analysis, a total set of 11 groups of replicates (at least two samples) passed the filtering criteria (over 10,000 reads). The PCoA visualization at genus and species level showed a dispersion of the different samples, but with a clustering according to the respective replicates (Figure 4). This suggests that there is limited within-sample variation when the same samples are processed on different days by different operators.

### Relative abundance of 32 bacterial targets in healthy vaginal samples

To determine healthy reference ranges for the bacterial targets in the assay, we selected a set of 50 samples from our database. These represent self-reported healthy individuals from the uBiome microbiome research study. In addition to health status, additional selection criteria included no usage of antibiotics six months prior, and no current urinary tract or vaginal infections, including the presence of STDs. The 50 samples were processed with the target database, and the relative abundance ranges for each bacterial target in the cohort are shown (Figure 5).

**Figure 5.**
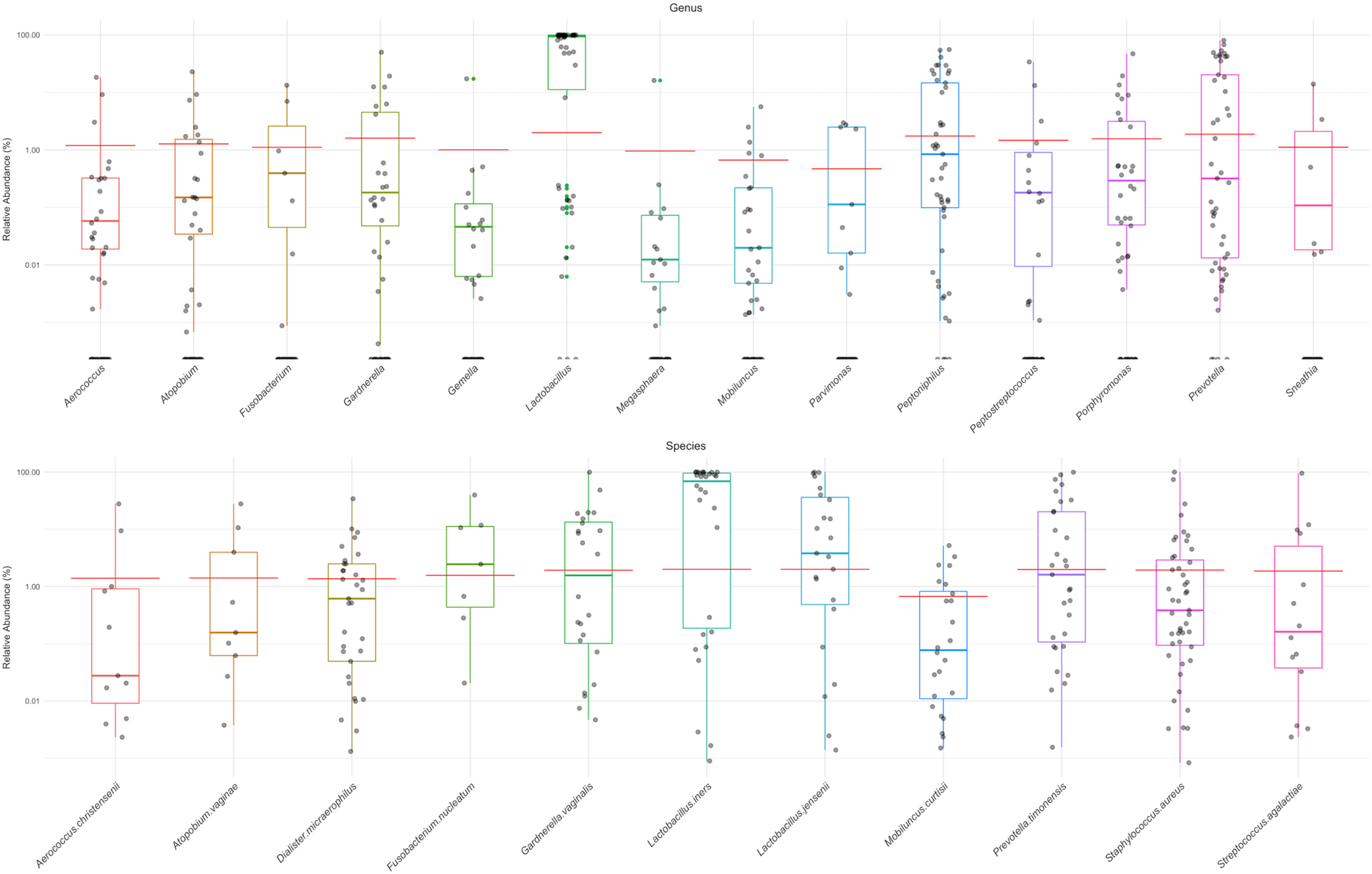
**Healthy ranges for the bacterial targets in the assay.** A set of 50 vaginal samples, each from a different woman, was selected based on the self-reported answers given to survey questions indicating general and vaginal health. Each dot represents the relative abundance of a different bacterial target on genus level (top) or species level (bottom) within a different vaginal sample. Boxes indicate the 25th-76th percentile, with the median indicated inside each colored box. Red line indicates the 99% confidence interval of each distribution. Not all of the taxa used in the assay were plotted, as some had no abundance values for this healthy cohort (*Papillibacter*, *C. trachomatis*, *M. mulieris*, *M. genitalium*, *N. gonorrhoeae*, *P. amnii*, *T. pallidum*), based on the exclusion criteria.

As expected, given the nature of the samples, *Lactobacillus* was the most abundant genus, with the widest abundance distribution. At the species level, a similar distribution of the relative abundances was found, including a wide range and a high relative abundance for *Lactobacillus iners*.

### Pathogen detection

Among the 32 bacterial targets in the assay are four pathogens implicated in STI: *C. trachomatis*, *N. gonorrhoeae*, *M. genitalium*, and *T. pallidum*. The performance of the assay to detect two of these pathogens was confirmed on a set of ten clinical samples available through a commercial source, five of which were positive for *C. trachomatis*, and five of which were positive for *N. gonorrhoeae*. A vaginal pool consisting of samples derived from 11 healthy individuals was included as a control sample, and was found to be negative (Figure 6).

**Figure 6.**
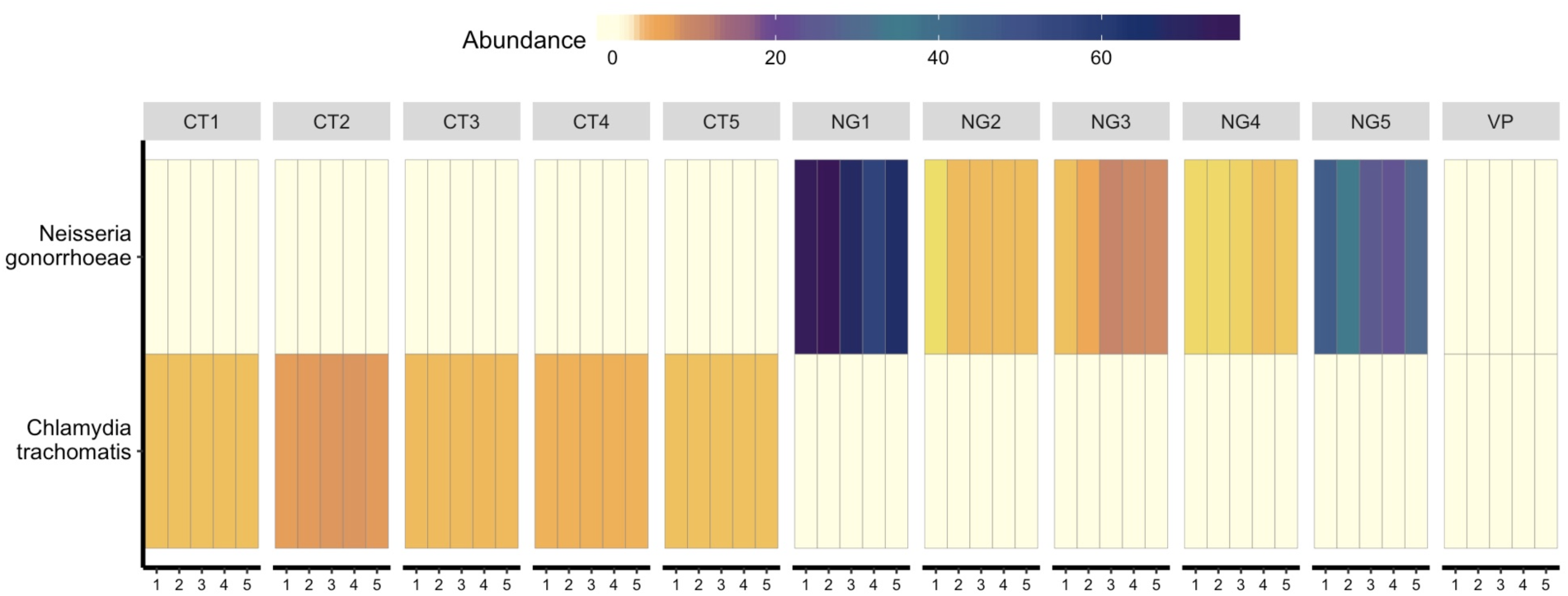
**Experimental validation of 16S rRNA gene sequencing for pathogen detection using verification samples.** Ten verification specimens (iSpecimen) containing either *C. trachomatis* (n=5) or *N gonorrhoeae* (n=5), as well as a vaginal pool (VP) constructed by combining 96 vaginal samples from 11 individuals, were tested for the presence of either pathogen using 16S rRNA gene amplification and sequencing. Five replicates of each specimen were tested. The heatmap shows the relative abundance of the two pathogens in each replicate experiment, on a scale from light yellow (absent) to dark blue (100% relative abundance).

The four STI-associated targets (*C. trachomatis*, *M. genitalium*, *N. gonorrhoeae*, and *T. pallidum*) were not present in any of the 50 samples from the healthy subject set (see also below), nor in a set of 88 vaginal samples used to validate the performance of the Digene test on extracted DNA (see below). In a set of 185 samples used to compare the HPV genotyping part of the assay to the Digene test (see below), *C. trachomatis* was present in one of the samples, while the other STI targets were not found.

### Performance of Digene HC2 HPV test on extracted DNA

In order to validate the use of the Digene kit on extracted vaginal DNA, we compared the performance of the Digene HC2 HPV assay on a set of 88 self-obtained, paired vaginal samples, i.e. a Digene brush resuspended in Digene STM, as well as DNA extracted from a paired vaginal swab resuspended in lysis transport medium. Of the 88 samples, 85 showed concordant results (70 were negative and 15 were positive for HPV in both tests) (Table 1). Three sample pairs that were positive with the Digene brush were HPV negative when the corresponding test was performed on extracted DNA. These three samples had an average Digene RLU ratio of 1.94, suggesting that these contained low levels of HPV (Figure 7).

**Table 1:** **Digene HC2 High-Risk HPV assay performance on a set of 88 paired, self-collected vaginal samples.** One set of samples was collected using a Digene brush resuspended in Digene Specimen Transport medium (“Digene STM”), and the second set was extracted DNA from swabs suspended in tubes with lysis/stabilization buffer (“Digene DNA”). Samples were considered to be hrHPV positive if the RLU ratio was 1 or more. Agreement between Digene STM and Digene DNA was of 96.6%, while concordance by Cohen’s Kappa was 0.89 ± 0.12 (Z=8.39, p-value=0.0001).

|  | Digene DNA<br>hrHPV - | Digene DNA<br>hrHPV + | Sum |
| --- | --- | --- | --- |
| Digene STM hrHPV - | 70 | 0 | 70 |
| Digene STM hrHPV + | 3 | 15 | 18 |
| Sum | 73 | 15 | 88 |

**Figure 7.**
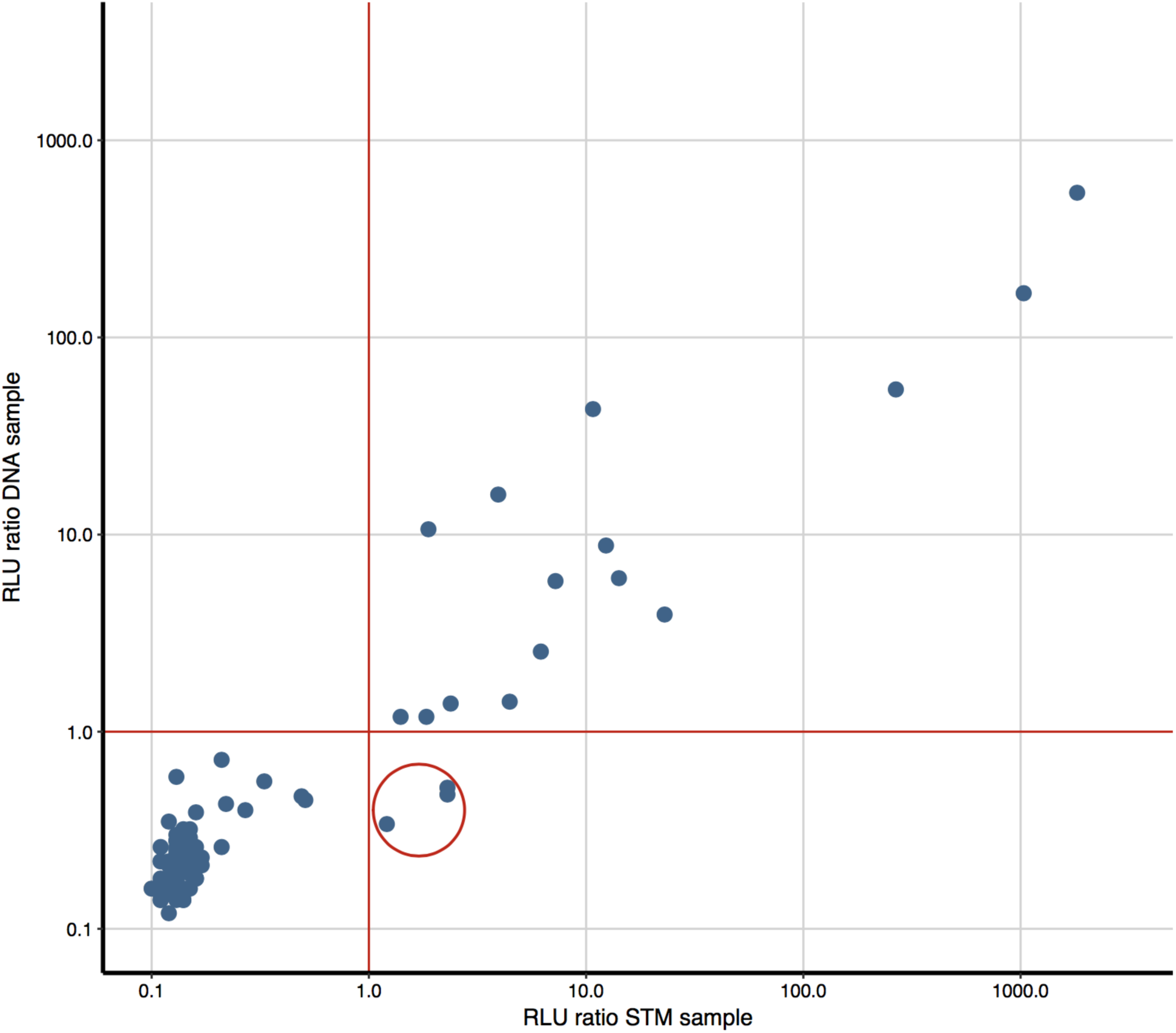
**Digene HC2 High-Risk HPV assay performance on a set of 88 paired, self-collected vaginal samples**. Samples were tested directly from STM tubes (X axis) or from a paired sample after DNA extraction (Y axis). The red lines show the cutoff of the Digene assay (RLU ratio = 1). The red circle highlights three STM specimens that were positive for hrHPV with an average RLU ratio of <2 (low positive), but below RLU ratio=1 for their corresponding extracted DNA specimen. The results for all other 85 specimens were concordant.

### Performance of the HPV sequencing test on clinical samples

Using 185 vaginal specimens, the performance of the assay to detect hrHPV was compared to that of the Digene HC2 hrHPV assay. The Digene test was considered positive if the measured RLU was equal to or greater than the assay’s cutoff (RLU ratio of 1 or higher), as per the manufacturer’s instructions. The assay was considered to be positive for hrHPV if the normalized number of reads assigned to hrHPV types divided by the number of reads assigned to a spiked-in control was greater than 0.1. Of the 185 samples, 145 were negative in both tests, while 36 were positive in both tests (Table 2), with an overall agreement of 97.83%, Cohen’s kappa = 0.93 ± 0.064. Two samples were positive in the Digene HC2 hrHPV assay but did not yield any hrHPV reads after amplification and sequencing by our assay. Of these two false negatives, however, one contained lrHPV 61, and the other one contained both lrHPV 30 and lrHPV 61. Although these two lrHPV types were not validated for our assay, their sequences were identified after amplification and genotyping. Two false positive samples were negative in the Digene HC2 hrHPV assay but yielded sufficient hrHPV reads to be identified as positives by our genotyping assay. One of these samples contained hrHPV type 35, and the other contained hrHPV type 68. Thus, in comparison to the Digene HC2 hrHPV test, the hrHPV sequencing assay had a sensitivity of 94.74% and a specificity of 98.64%.

**Table 2.** **Comparison of the women’s health assay for the detection of genotyped hrHPV in DNA extracted from 185 vaginal samples to that of the Digene HC2 hrHPV assay.** Concordance by Cohen’s kappa (0.93 ± 0.064, Z = 12.7, p-value=0.0001), shows that the two test are in excellent agreement.

|  | Genotyping<br>hrHPV - | Genotyping<br>hrHPV + | Sum |
| --- | --- | --- | --- |
| Digene hrHPV - | 145 | 2 | 147 |
| Digene hrHPV + | 2 | 36 | 38 |
| Sum | 147 | 38 | 185 |

Excellent correlation was found between the number of normalized hrHPV sequencing reads and the Digene HC2 hrHPV RLU ratios, confirming that the PCR and sequencing hrHPV assay described here can not only detect hrHPV types but also assess their relative abundance (Figure 8).

**Figure 8.**
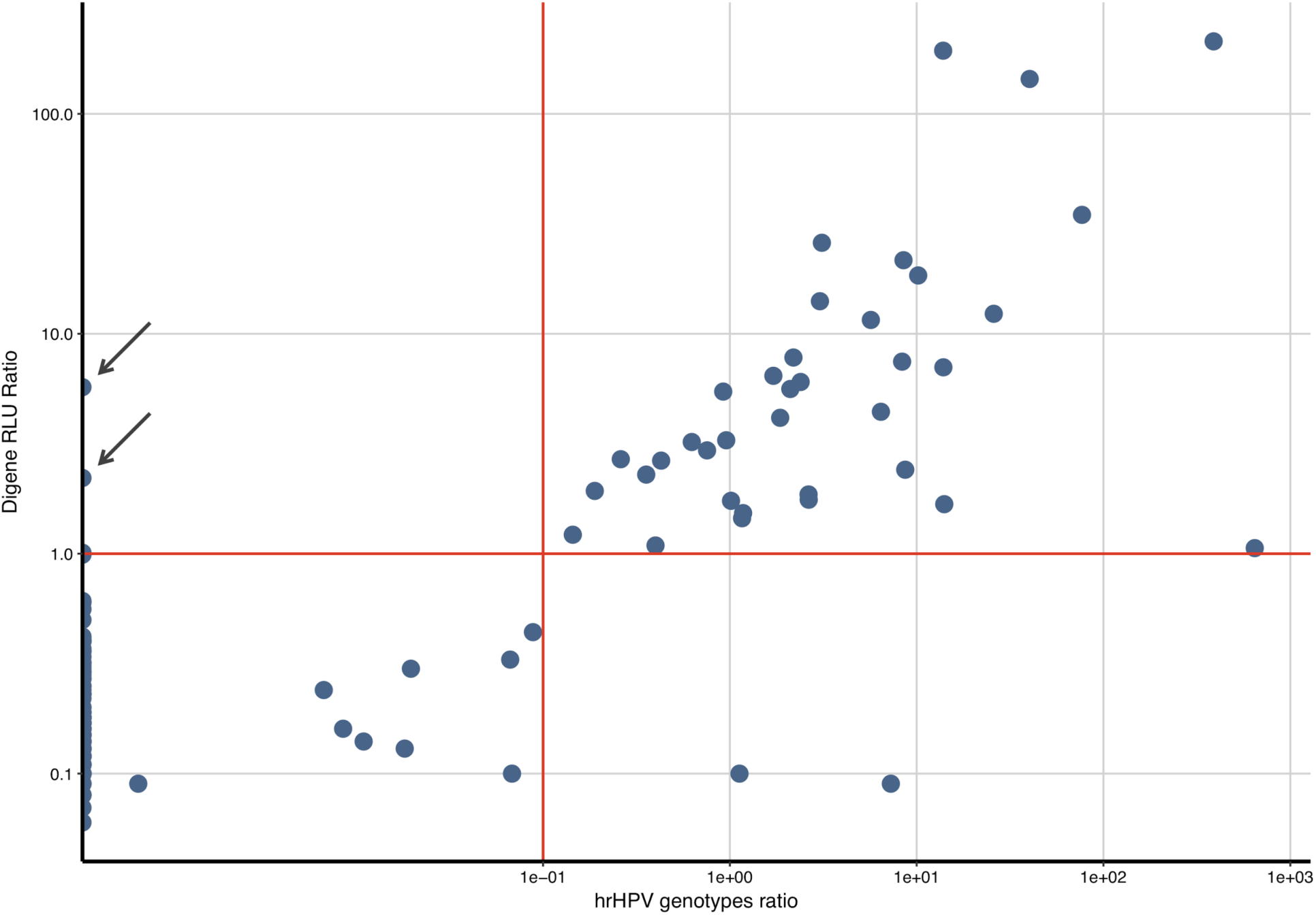
**Comparison of hrHPV amplification and sequencing to the Digene HC2 hrHPV assay.** DNA from 185 vaginal samples was extracted and tested by PCR amplification and sequencing using HPV primers, and additionally used directly in the Digene assay using the HC2 hrHPV probe mix. For each sample, X-axis shows the normalized ratio of reads assigned to hrHPV to reads assigned to a spiked-in internal control, while the Y-axis shows the Digene hrHPV probe RLU values normalized over the assay’s cut-off RLU. The red lines show the cutoff for each of the assays. Two samples that were positive in the Digene hrHPV assay but in which no hrHPV sequences could be detected are pointed out with arrows.

## Discussion

Here, we describe a novel women’s health assay combining vaginal microbiome analysis, STI-associated pathogen detection, and HPV detection and identification in a self-sampling format. Although each of these components have been described before, to our knowledge, this assay is the first to combine all of these parts, thus offering women a unique opportunity to gain a broad perspective into their vaginal and reproductive health.

The detection of hrHPV in combination with self-sampling has been proposed as an effective method for cervical cancer risk screening. Although the sensitivity of signal-based hrHPV detection, such as the Digene assay, in self-obtained vaginal swabs has been found to be slightly lower than that in clinician-obtained cervical specimens, hrHPV detection based on PCR was shown to be equally sensitive in self-sampled specimens (Arbyn et al. 2014). While this assay is not intended to replace regular cervical cancer screening programs, offering women the opportunity to self-collect vaginal specimens poses fewer barriers for women to be screened, and thus could lead to increased participation rates. Therefore, encouraging women to self-collect vaginal samples for hrHPV screening, as already implemented in numerous countries, and recommending them to seek further physician examination in case of a positive result, while encouraging them to participate in regular screening programs may have a positive impact on rates of detection of cervical cancer, and potentially save lives (Wong et al. 2016).

Unlike most currently available clinical assays, this assay not only detects whether hrHPV is present in a sample, but also identifies the presence of specific type(s) by using sequencing analysis. This is of particular importance because the prevalence of hrHPV types might shift in the setting of recently introduced HPV vaccines. In the last decade, many countries, including the US, have implemented HPV vaccination programs (Harper and DeMars 2017) and the prevalence among young women of hrHPV types covered by these vaccines such as HPV types 16 and 18 is rapidly dropping (Drolet et al. 2015). It is too early to know if other HPV types might become more prevalent, but sequence-based HPV detection offers the ability to detect types that are not covered by tests that focus on the detection of a limited number of types.

Several other HPV genotyping assays based on PCR and sequencing have been reported (Gradíssimo and Burk 2017). Oliveira Carvalho and coworkers showed that their PCR and sequencing-based method was 4-fold more likely to identify the viral type in hrHPV positive samples than type-specific PCR (Carvalho et al. 2010). Another study showed that the use of sequencing detected HPV types not found by traditional methods, with a detection specificity of 100% in comparison with PCR and p16 immunohistochemistry (Conway et al. 2012). Using 454 sequencing of PCR amplicons and artificial and clinical sample sets, Militello et al. found that HPV sequencing performed well and was capable to detect infections with multiple types and even detect novel types, compared to other assays including the Digene HC2 test (Militello et al. 2013). A more recent study compared the use of ion-torrent sequencing with the Linear Array (LA) method for genotyping of anal hrHPV (Nowak et al. 2017). This study showed that the sequencing method was accurate and able to detect variants that LA did not, and that it could detect multiple types in multiple HPV infections. Ambulos et al. showed high sensitivity of a sequencing-based HPV genotyping assay performed on formalin-fixed paraffin-embedded oropharyngeal and cervical specimens from subjects with cancer; this method showed high concordance (92%) in comparison with LA-based genotyping (Ambulos et al. 2016). Thus, HPV detection by PCR and sequencing shows excellent performance in comparison to currently used clinical assays.

As recommended by the VALGENT study framework (Arbyn et al. 2016), we compared the performance of the HPV component of the test to that of the widely used Digene HC2 hrHPV assay. Because the novel test reported here is performed on extracted DNA, we first validated the use of the Digene assay on extracted DNA. The Digene performance on the extracted DNA was slightly less sensitive than that directly performed on the Digene STM tubes; 3 out of 18 samples that tested positive in the Digene assay on STM tubes subsequently tested negative on their corresponding extracted DNA. The Digene HC2 assay has been found to give discordant results in about 8% of paired tests (Carozzi et al. 2005; Castle et al. 2002), where, for example, a positive sample will test negative at retesting, and the majority of those samples will have a low RLU ratio between 1.00 and 3.00 in the positive test. Reproducibility testing using the Digene assay therefore is expected to be lowest in samples with an RLU ratio near the cutoff value, and a cutoff ratio of 2 or 3 instead of 1 has been proposed to serve as a better indicator for reproducible positive results (Carozzi et al. 2005, Castle et al. 2002, de Cremoux et al. 2003, Moss et al. 2015). In our study, all specimens with RLU ratios of 2 or higher in the direct Digene test on STM tubes were also correctly identified as positive when the test was performed on extracted DNA, suggesting that the Digene assay can be applied to extracted DNA as well.

Using extracted DNA from 185 vaginal specimens as the template, the performance of the hrHPV sequencing assay was compared to that of the Digene HC2 hrHPV assay. The hrHPV sequencing assay had excellent correlation with the Digene assay, with a sensitivity and specificity of 94.74% and 98.64%, respectively, and a Cohen’s kappa of 0.93. In addition, the two tests were in good general agreement about the relative amount of HPV molecules detected. Of the two samples that were reported positive by the Digene assay but that did not contain hrHPV sequences as determined by our assay, both of them were found to contain lrHPV types. Cross-reactivity of the Digene HC2 hrHPV probe mix with lrHPV sequences such as 30 and 61 has been demonstrated by several others (Boehmer et al. 2014; de Cremoux et al. 2003; Gillio-Tos et al. 2013; Ginocchio et al. 2008; Vernon et al. 2000). Thus, even though these samples had to be classified as false-negative because the Digene assay was taken as the gold standard, it is likely they were actually false-positives in the Digene test.

In addition to the HPV portion of the novel women’s health assay described here, the assay also reports the relative abundance of commensal and pathogenic bacteria in vaginal samples. Self-collection has been shown to be well-suited for vaginal microbiome analysis as reported by Forney et al. who showed that microbial diversity is similar between self-collected and physician collected vaginal samples (Forney 2010).

Several bacteria have been associated with vaginal health conditions, such as bacterial vaginosis (Ling et al. 2010, Ravel et al. 2011, Ravel and Wommack 2014, Srinivasan et al. 2012), aerobic vaginitis (Donders et al. 2017), pelvic inflammatory disease (Brunham et al. 2015), and sexually transmitted infections (Hill et al. 2016; Jensen 2017; Petrova et al. 2015, Ziklo et al. 2016). The women’s health assay described here detects the relative abundance of bacteria positively associated with bacterial vaginosis, such as *Sneathia* or *Gardnerella* species, as well as those negatively associated with that condition such as *Lactobacillus* species. In addition, it detects the presence of four common STI-associated pathogens, i.e., *C. trachomatis*, *N. gonorrhoeae*, *M. genitalium*, and *T. pallidum*. Of these, *M. genitalium* has been recently recognized as an important pathogen implicated in pelvic inflammatory disease and infertility (Jensen 2017; Wiesenfeld and Manhart 2017). Although some early diagnostic tests have been described (Gaydos 2017, Munson 2017), very few clinicians test for its presence. Furthermore, the vaginal microbiota composition has been reported to be associated with the progression of HPV infection, from early states to cervical cancer (Brotman 2014, Mitra et al. 2016). Vaginal microbiome analysis therefore not only can be used to detect STI-associated pathogens and bacteria involved in bacterial vaginosis, but also to assess a woman’s microbiome similarity to the microbiome of a group of individuals with progressed HPV infection. This brings about the opportunity to leverage microbiome information to understand HPV infection progression and women’s susceptibility to cancer development.

In conclusion, we here present a women’s health assay that for the first time combines the detection of the most important bacterial and viral indicators of vaginal health and disease. We envision that this test will greatly help women to learn about their vaginal microbiome, encourage them to participate in existing cervical screening programs because it allows for self-sampling, and assist their doctors to more accurately diagnose and treat diseases of the genital tract.

